# Integrative Multi-Scale Sequence–Structure Modeling for Antimicrobial Peptide Prediction and Design

**DOI:** 10.64898/2026.02.24.707610

**Authors:** Jiayi Li, Yanruisheng Shao, Yu Li, Qinze Yu

## Abstract

Antimicrobial resistance (AMR) is accelerating worldwide, undermining frontline antibiotics and making the need for novel agents more urgent than ever. Antimicrobial peptides (AMPs) are promising therapeutics against multidrug-resistant pathogens, as they are less prone to inducing resistance. However, current AMP prediction approaches often treat sequence and structure in isolation and at a single scale, leading to mediocre performance. Here, we propose MultiAMP, a framework that integrates multi-level information for predicting AMPs. The model captures evolutionary and contextual information from sequences alongside global and fine-grained information from structures, synergistically combining these features to enhance predictive power. MultiAMP achieves state-of-the-art performance, outperforming existing methods by over 10% in MCC when identifying distant AMPs sharing less than 40% sequence identity with known AMPs. To discover novel AMPs, we applied MultiAMP to marine organism data, discovering 484 high-confidence peptides with sequences that are highly divergent from known AMPs. Notably, MultiAMP accurately recognizes various structural types of peptides. In addition, our approach reveals functional patterns of AMPs, providing interpretable insights into their mechanisms. Building on these findings, we employed a gradient-based strategy and achieved the design of AMPs with specific motifs. We believe MultiAMP empowers both the rational discovery and mechanistic understanding of AMPs, facilitating future experimental validation and therapeutic design. The codebase is available at https://github.com/jiayili11/multi-amp.

## 1 Introduction

Antimicrobial resistance (AMR) is among the most critical public health challenges today due to the overuse of antibiotics. AMR is estimated to cause up to 10 million deaths annually by 2050 [1], and it requires urgent measures. However, the development and approval of new antibiotics have declined in recent decades. Therefore, alternative therapeutic approaches are essential to alleviate AMR and ensure the effective treatment of infectious diseases. Antimicrobial peptides (AMPs), which are key components of the innate immune system in animals [2], are regarded as one of the most promising alternatives to antibiotics. They offer effective control of drug-resistant pathogens while minimizing the risk of resistance development. For example, the human cathelicidin LL-37 acts as a cornerstone of our innate immunity by directly killing microbes and modulating the host’s immune response [3]. Nisin, produced by bacteria, showcases a highly specific mechanism that has led to its successful commercial use as a natural food preservative [4]. The synthetic peptide SAAP-148, an optimized derivative of a human protein, was specifically developed to combat the urgent threats posed by multidrug-resistant bacteria and their notoriously resilient biofilms [5].

In recent years, AMPs have attracted considerable research interest, leading to the development of computational approaches to accelerate their discovery and design [6, 7, 8, 9, 10, 11, 12]. Conventional methods for AMP prediction typically rely on learning from existing peptide sequences. AmPEP [13] takes advantage of distribution patterns of amino acids, AMPScanner [14] directly uses peptide sequence as input and leverages the power of CNN and LSTM, AMP-BERT [15] and PepNet [16] employ pre-trained protein language models to enrich sequence representation. These frameworks exhibit promising performance on in-distribution data. However, known AMPs represent only a small portion of the vast peptide sequence space, meaning that many unrevealed AMPs with various sequences remain undiscovered. Our recent research [17] demonstrates that, despite the sequence diversity of AMPs, they predominantly adopt canonical structural folds such as *α*-helical, extended, and *β*-sheet conformations, suggesting AMPs converge toward effective antimicrobial architectures. Hence, one strategy to handle the out-of-domain peptides is to incorporate structural information, which has been broadly applied in protein-related tasks. GS-DTI [18] extracts the contact map from the tertiary structure to model the protein graph for drug-target interaction prediction, while graphEC [19] takes the distance, direction, and orientation features from the structure to construct the geometric graph for protein representation. Moreover, recent studies [20, 21, 22, 23] have demonstrated that integrating multi-modal information that includes both sequence-based and structure-based features can further promote model performance. Inspired by those, we proposed MultiAMP, an AMP prediction framework that exploits peptide representations from different scales and modalities, including sequence contextual information, evolutionary information, global secondary structure, and detailed tertiary structure, to enhance the performance on predicting remote AMPs that share low sequence identity with training data (Figure 1A). Our approach takes raw amino acid sequences as input and jointly predicts AMP activity and secondary structure. The integration of secondary structure prediction enables the model to capture conserved structural motifs that are often essential for peptide function. This structural awareness boosts the generalization ability of the model, particularly in detecting distant AMPs that might be overlooked by sequence-based approaches alone. We compared MultiAMP with seven task-related models, and our framework out-performs all the previous methods by a large margin on various evaluation criteria in the 40% sequence similarity cut-off test. Through the embedding analysis, we found that MultiAMP encodes information about distinctions between AMPs and non-AMPs, as well as the structural types of peptides. We then applied the model to marine organism data to discover novel AMPs, identifying 484 high-confidence candidates with diverse sequences. Despite their low sequence identity to previously characterized AMPs, the majority adopt conventional *α*-helical structures. Interpretable analysis of our model demonstrates its ability to accurately identify functional motifs in novel AMPs, which are predominantly cationic and amphipathic and characterized by a high frequency of basic residues such as lysine (K) and arginine (R)—hallmark features of antimicrobial peptides. Based on these findings, MultiAMP is adopted for motif-constrained AMP design. Generated peptides preserved intended structural features while exhibiting improved antimicrobial activity and low toxicity (Figure 1B). Our method enables accurate and interpretable prediction and design of AMPs, offering significant potential for applications in precision medicine.

**Figure 1:**
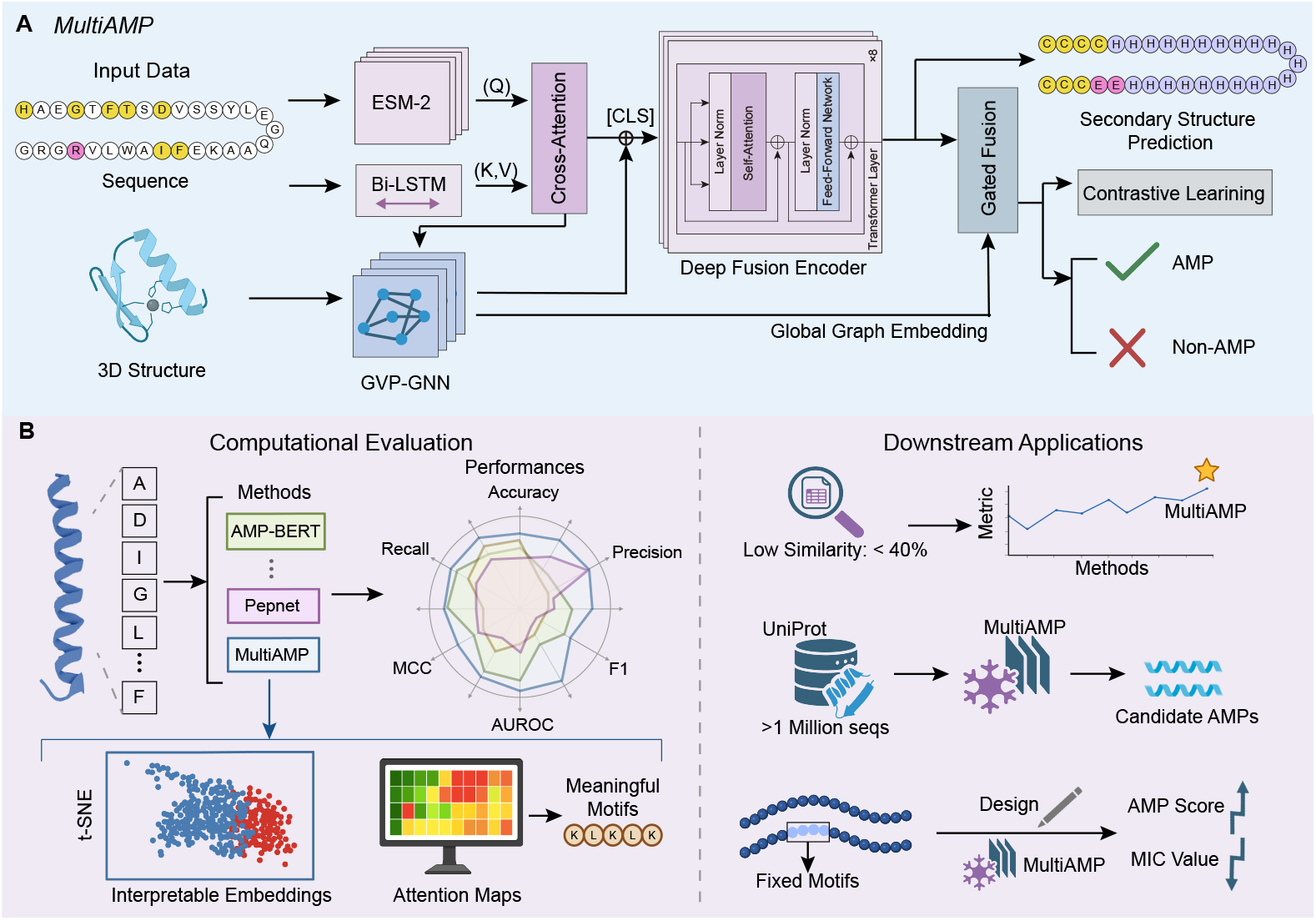
MultiAMP integrates multi-level and -modal learning with structure-aware encoding for comprehensive AMP analysis. **(A)** Architecture overview. ESM-2 and Bi-LSTM for sequence features, GVP-GNN for geometric encoding. Cross-attention integrates both modalities, feeding into a Deep Fusion Encoder with gated fusion for final classification. Multi-task learning jointly predicts AMP activity and secondary structure (3-state), while contrastive learning enhances discrimination. **(B)** Evaluation and applications. Left: Benchmark comparison against 7 methods across multiple metrics. We utilize t-SNE embeddings to show clear AMP/non-AMP separation and attention maps to highlight functional motifs. Right: Three applications: (1) performance verification on remote AMPs; (2) mining AMPs from ocean data; (3) de novo AMP design with specific motifs.

## 2 Methods

### 2.1 Data Construction

#### Training data

To eliminate potential bias arising from differences in sequence length between positive and negative samples, the training dataset was constructed to maintain both length distribution and quantity balance across the two groups. The positive dataset was compiled from the DBAASP database [24], with CD-HIT [25] applied at a 90% similarity threshold, resulting in 5,985 AMPs. For negative samples, we queried UniProt [26] with exclusion criteria for antimicrobial-related annotations, then paired each AMP with a length-matched non-AMP, yielding 5,985 negative samples with identical length distributions.

#### Test data

To ensure rigorous evaluation of generalizability, we curated a comprehensive test set by integrating four distinct sub-datasets orthogonal to the training set [27, 6]. After removing duplicates and overlapping sequences, the final test set comprised 1,234 positive and 4,121 negative samples. The specific distribution comparison between training and test sets is illustrated in Supplementary Figure **1**.

#### Peptide structure

Secondary structures were predicted by PHAT [28], and tertiary structures were obtained using ESMFold [29].

### 2.2 Model Framework

MultiAMP integrates three distinct information streams to generate comprehensive peptide representations for AMP classification. Specifically, it employs ESM-2 [29], a pre-trained protein language model, to encode evolutionary and functional features; a bidirectional LSTM (BiLSTM) module to capture contextual sequence interactions; and Geometric Vector Perceptrons Graph Neural Networks (GVP-GNN) [30] to model fine-grained geometric and structural information. To further enrich representations, we incorporate a secondary structure reconstruction task into our framework. In addition to AMP classification, MultiAMP is trained to simultaneously predict the secondary structure of peptide sequences. This strategy introduces a strong inductive bias, encouraging the model to identify sequence patterns and structural motifs associated with secondary structure formation, which are often linked to antimicrobial activity. As a result, the learned embeddings are endowed with biologically relevant information, ultimately enhancing the robustness and accuracy of AMP classification.

#### Multi-scale feature extraction

MultiAMP extracts complementary information from three sources: (1) Evolutionary and contextual features via ESM-2, aggregating representations from layers ℒ = { 6, 12, 18, 24, 33 } using learnable weights *w*_*l*_, normalized via softmax, to capture multi-scale evolutionary patterns:

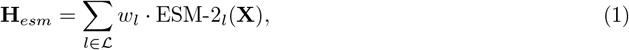

where **X** represents the tokenized protein sequence and the weights are jointly optimized during training; (2) Sequential dependencies via BiLSTM (hidden size 128 per direction) to model local context and position-specific patterns, producing contextualized embeddings **H***raw* ∈ R^*L*×256^; (3) for Geometric structural features, we apply GVP-GNN with 3 layers, 384 scalar dimensions, and 4 vector dimensions. This module processes 3D coordinates to encode backbone geometry, dihedral angles, and spatial relationships within 10Å neighborhoods. The GVP-GNN takes initial scalar features (amino acid identity, backbone dihedral angles *ϕ, ψ, ω*, position embeddings) and vector features (backbone unit vectors, normal vectors, tangent vectors) as input, then performs rotation-equivariant message passing through:

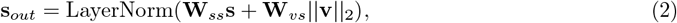

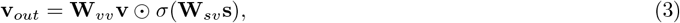

where **s** and **v** denote scalar and vector features, **W***ss*, **W***vs*, **W***vv*, and **W***sv* are learnable weight matrices, and *σ*(·) is the SiLU activation. Detailed feature specifications are provided in Supplementary Note **1**.

#### Hierarchical multi-modal fusion

Integration of sequence and structural modalities proceeds through two stages. In the first stage, Cross-attention alignment is performed between ESM-2 evolutionary features, which serve as queries, and BiLSTM contextual features, which serve as keys and values. This 8-head multi-head attention mechanism includes a residual connection:

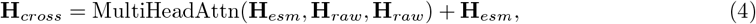

enabling the model to refine pre-trained representations with explicit sequential context. In the second stage, Transformer-based deep fusion processes the concatenated cross-attended sequence features and GVP-GNN structural embeddings **H***struct* through 3 transformer encoder layers with 8 attention heads and a hidden dimension of 512:

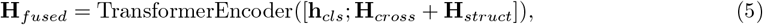

where **h**_*cls*_ is the CLS token from ESM-2. The final representation combines sequence-level features, specifically the CLS token from the transformer output denoted as 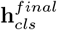, with graph-level features through a gated fusion mechanism:

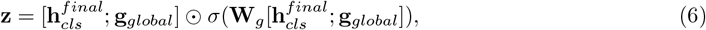

where **W**_*g*_ is a learnable gating matrix and *σ*(·) is the sigmoid activation. This gating mechanism adaptively balances modality contributions based on input characteristics, as visualized in Figure 3C.

#### Multi-task learning framework

MultiAMP employs a multi-task learning strategy that jointly optimizes three complementary objectives: (1) primary binary classification for AMP prediction using Binary Cross-Entropy (BCE) loss; (2) supervised contrastive learning to structure the embedding space, encouraging same-class samples to cluster while pushing different-class samples apart:

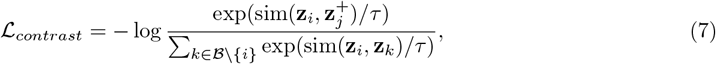

where sim(·,·) denotes cosine similarity, 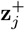 is a positive sample with the same label as anchor **z**_*i*_, ℬ is the training batch (balanced with equal AMPs and non-AMPs), *k* indexes all samples except the anchor, and *τ* = 0.07 is the temperature; and (3) secondary structure reconstruction guided by a composite loss combining focal loss (addressing class imbalance with focusing parameter *γ* = 3.0 and class-balancing weights *α*_*t*_), Conditional Random Field (CRF) loss (capturing position dependencies via a learned 3 × 3 transition matrix for H/E/C states), and continuity regularization (promoting contiguous structural segments):

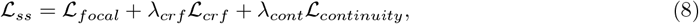

where λ _*crf*_ = 0.3 and λ _*cont*_ = 0.1. The continuity term penalizes abrupt transitions between adjacent predictions:

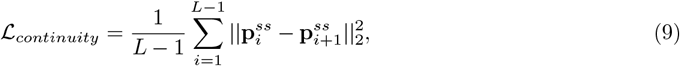

where 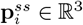 is the predicted probability distribution over three secondary structure states at position *i*. The total objective combines these components with dynamic weights:

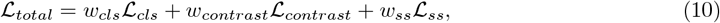

where *w*_*cls*_ = 1.0 (fixed), and auxiliary task weights (*w*_*contrast*_ = 0.1, *w*_*ss*_ = 0.15, selected via grid search on the validation set) decay by 0.90 per epoch starting from epoch 25 to gradually emphasize the primary classification task. Detailed loss formulations and training configurations are provided in Supplementary Note **1**.

## 3 Results

### 3.1 MultiAMP exhibits superior generalization performance on remote antimicrobial peptides

We first assess the performance of our model alongside the other 7 AMP predictors, including iAMP-2L [31], AmPEP [13], AMAP [32], Macrel [33], AMPScanner Vr2 [14], AMP-BERT [15], and PepNet [16], on the curated test data that contains 5,355 sequences. Figure 2A illustrates the distribution of maximum sequence similarities between the test set and the training set, and the majority of test sequences are distinct from the training data. To ensure a fair comparison, all the models are retrained using the same training data as ours. Six evaluation metrics, namely Accuracy, Precision, Recall, F1-score, MCC, and AUROC, are applied for performance estimation. As illustrated in Figure 2B and C, MultiAMP consistently outperformed all baselines, achieving an AUROC of 0.9810, which is substantially higher than the next-best model, PepNet (AUROC = 0.9642). Similarly, MultiAMP attained the highest average precision (AP = 0.9560), as shown by the precision-recall curves. Figure 2D and Supplementary Table **1** further demonstrate that MultiAMP yields the highest scores across all metrics, with an MCC of 0.8519, an F1-score of 0.8857, a Precision of 0.8923, and an Accuracy of 0.9477. These results underscore the superior classification capability of MultiAMP compared to established methods, especially in distinguishing AMPs from non-AMPs within diverse sequence backgrounds.

**Figure 2:**
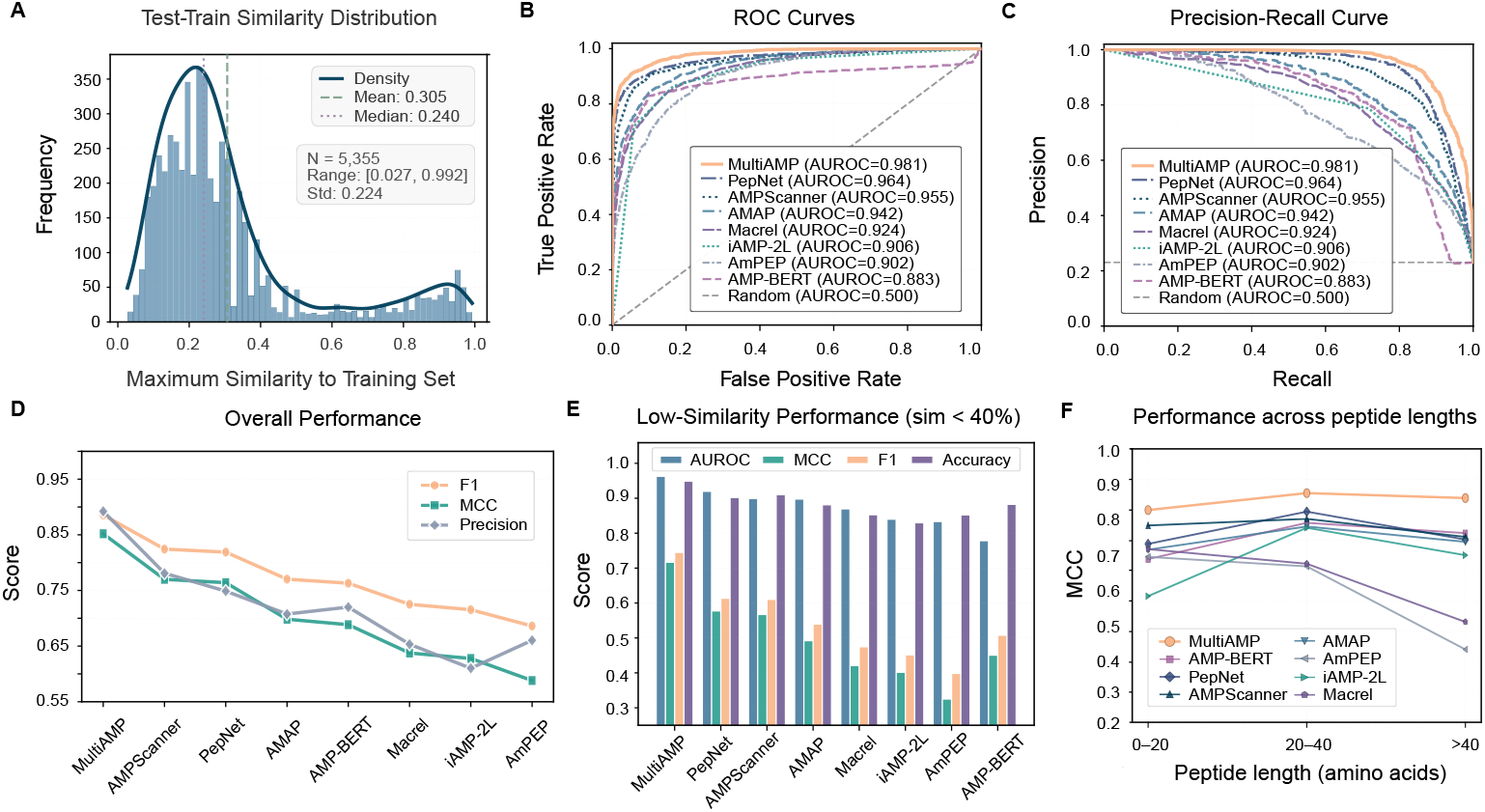
MultiAMP achieves superior performance and generalization through multi-level architecture integration. **(A)** Sequence similarity distribution between the test data and their closest train data. **(B)** ROC curves comparing MultiAMP against 7 baselines and random case on 5,355 test peptides. **(C)** Precision-Recall curves for all methods. Baseline represents random performance. **(D)** Overall performance on the test set across F1, MCC, and Precision metrics. **(E)** Model performance on test data with low sequence similarity to the training set (< 40% identity). **(F)** Performance benchmark testing of models under different peptide length groups (0–20, 20–40, > 40 aa).

A crucial requirement for practical AMP prediction is the ability to generalize to novel peptides, particularly those with low sequence identity to the training set. To rigorously evaluate this, we filtered the test set to exclude any sequences sharing more than 40% identity with the training data, resulting in a challenging subset of 4,379 peptides. As depicted in Figure 2E and Supplementary Table **1**, all baseline models suffered marked drops in performance, with F1-scores declining by at least 0.2. In stark contrast, MultiAMP maintained strong generalization, achieving an F1-score of 0.7451 and an MCC of 0.7169, outperforming all comparators by significant margins (minimum F1-score improvement: 0.13 over PepNet). Notably, MultiAMP also led in AUROC (0.9629) and accuracy (0.9491) in this low-similarity regime. These results reflect its robust capacity to recognize truly novel AMPs beyond sequence memorization. For completeness, we also assessed performance on the subset of peptides exhibiting ≥ 40% sequence identity with the training data (Supplementary Table **1**). Here, as expected, most models showed improved metrics, and MultiAMP continued to lead or match the top scores.

In addition to sequence similarity, we also evaluated performance with respect to peptide length. The test data were divided into three groups based on sequence length: less than 20, between 20 and 40, and greater than 40 residues. As shown in Figure 2F, MultiAMP outperforms all comparative methods across all peptide length groups, and keeps the MCC value above 0.8, demonstrating robust and stable performance regardless of peptide length. In contrast, the majority of baseline methods exhibited substantial variability in MCC across different length groups. For instance, while AMAP and AMPScanner performed comparably well for peptides in the 20–40 residue range, their performance declined for longer peptides. Other methods, such as AmPEP and iAMP-2L, showed even greater sensitivity to peptide length, with significantly reduced MCC values for longer peptides.

In summary, MultiAMP demonstrates state-of-the-art performance across various evaluated scenarios. These results collectively establish MultiAMP as a robust and trustworthy tool for AMP prediction, addressing key limitations of existing methods in both accuracy and generalizability.

### 3.2 Multi-level architecture integrates sequence and structural features for robust prediction

To systematically assess the contribution of each module to MultiAMP’s performance, we conducted comprehensive ablation experiments on both the complete test dataset and a subset comprising sequences with less than 40% identity (low similarity), as well as a high-similarity subset (≥ 40% identity). Six model variants were evaluated: (1) removal of the GVP-GNN structure encoder (w/o GVP); (2) withoust secondary structure prediction auxiliary task (w/o SS); (3) omission of both the GVP-GNN encoder and the secondary structure prediction auxiliary task (w/o GVP+SS); (4) exclusion of the BiLSTM module responsible for capturing local sequential context (w/o BiLSTM); (5) a baseline model utilizing only the ESM-2 encoder (w/o GVP+SS+BiLSTM); and (6) a variant without the ESM-2 encoder (w/o ESM2).

The ablation study (Supplementary Table **2**) shows that each architectural component of Multi-AMP is essential for optimal performance, particularly on the low-similarity (< 40% identity) test set, which evaluates the model’s generalization to novel or highly divergent sequences. The full architecture achieved the best results (AUROC = 0.9629, MCC = 0.7169, F1 = 0.7451). Removing either the GVP-GNN (AUROC = 0.9303, MCC = 0.6498) or ESM-2 (AUROC = 0.8914, MCC = 0.5397) led to sharp performance drops, confirming that integrating evolutionary and structural information is critical for robust generalization. The largest deficit occurred without ESM-2, highlighting the importance of deep evolutionary representations, while omitting GVP-GNN also reduced structural modeling capacity, as reflected by the secondary-structure prediction accuracy falling from 96.3% to 47.4%. In contrast, performance differences were minor on the high-similarity (≥ 40%) subset, where overlap with training data makes generalization less challenging. These results highlight that MultiAMP’s multi-modal design is particularly effective for the accurate prediction of novel AMPs, demonstrating superior generalization in discovering peptides with low evolutionary similarity.

### 3.3 Learned embeddings capture structural archetypes and functional motifs of antimicrobial peptides

To examine how MultiAMP captures peptide properties beyond sequence and structure patterns, we visualized our fused representations on test data using t-SNE. As shown in Figure 3A, AMP and non-AMP embeddings form clearly separated clusters, indicating that MultiAMP effectively encodes functional information.

**Figure 3:**
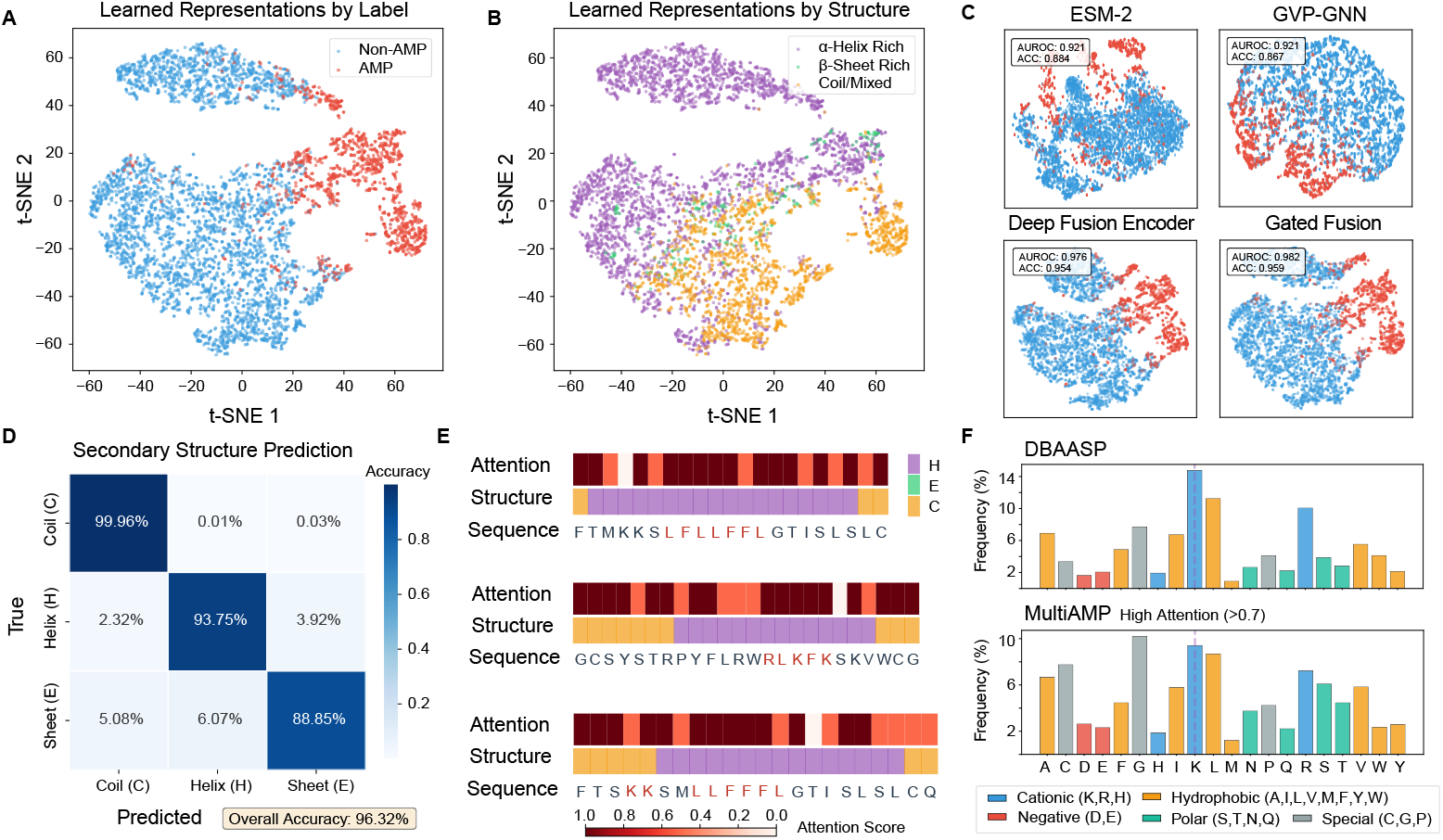
Learned representations of MultiAMP reveal structural organization and interpretable attention patterns. **(A)** Visualization of MultiAMP’s embeddings of test data using t-SNE. Datapoints are colored by labels. Clear AMP (red) and non-AMP (blue) separation indicates discriminative features. **(B)** The embeddings colored by secondary structure composition. Three structural archetypes emerge: helix-rich (purple), sheet-rich (green), and coil/mixed (orange). **(C)** T-SNE visualization of embeddings from different modalities. The upper panels display embeddings from individual modalities (ESM-2 and GVP-GNN), showing noticeable class overlap. The lower panels illustrate the effect of multi-modal fusion (Deep Fusion Encoder and Gated Fusion), which results in distinct class separation. **(D)** Confusion matrix for 3-state secondary structure prediction (helix, sheet, coil). **(E)** Attention weights analysis of MultiAMP on 3 samples. Regions with high attention scores overlaid with general AMP functional motifs. **(F)** Amino acid frequency in DBAASP database versus residues with high MultiAMP attention scores (> 0.7). The distributions are highly consistent (Pearson’s *r* = 0.822, *p* < 0.001), demonstrating that the model captures intrinsic AMP compositional patterns.

Grouping peptides by structural archetypes revealed additional organization (Figure 3B): *α*-helical peptides cluster tightly, while *β*-sheet and mixed/irregular peptides are more dispersed. These patterns suggest that MultiAMP learns biologically meaningful structural representations, particularly distinguishing *α*-helical peptides, which contributes to its strong predictive performance. To evaluate the impact of multi-modal integration, we visualized feature embeddings of different modules (Figure 3C). While individual modalities (ESM-2 and GVP-GNN) show moderate class separation, the fusion modules (Deep Fusion Encoder and Gated Fusion) produce significantly tighter and distinct clusters. This progression demonstrates that integrating sequence and structural data effectively refines the feature space, leading to superior classification performance.

Our multi-task framework yielded strong performance on secondary structure prediction, achieving an overall accuracy of 96.3% (Figure 3D), with class-specific accuracies of 93.75% for helices, 99.96% for coils, and 88.85% for sheets, indicating that the model effectively models structural features of peptides. For model interpretation, we analyzed the attention weights obtained from our Deep Fusion Encoder (Figure 3E). It revealed that the model prioritizes biologically meaningful sequence motifs and structural elements: for cationic AMPs, lysine and arginine clusters received maximal attention; for amphipathic helices, attention alternated between hydrophobic and charged faces, aligning with membrane-interaction models. Amino acid composition analysis (Figure 3F) further revealed enrichment of phenylalanine (F), lysine (K), and leucine (L) at high-attention sites, while glutamine(Q) and glutamic acid (E) are less frequent, confirming that MultiAMP selectively emphasizes residues critical for structure and function.

### 3.4 Systematic discovery reveals distinct antimicrobial peptides in ocean organisms

The discovery of novel antimicrobial peptides (AMPs) remains a critical priority to combat rising antimicrobial resistance and to broaden the chemical space of peptide therapeutics. However, current AMP repertoires are heavily biased toward a limited number of sequence families, constraining both generalization and rational design. To address this gap, we sought to identify previously uncharacterized antimicrobial peptides from ocean-derived proteins. Marine ecosystems exhibit exceptional phylogenetic and ecological diversity, making them a likely source of noncanonical AMP chemotypes. Their microbiomes span broad environmental gradients and experience intense biotic interactions that promote the evolution of membrane-active and metal-homeostasis peptides, positioning ocean datasets as a promising reservoir of structurally diverse AMPs [34, 35]. To systematically explore the antimicrobial potential of marine proteins, we constructed an ocean-focused peptide corpus by retrieving 104,693 sequences below 100 amino acids from UniProt [26] (Figure 4A). After excluding sequences with *≥* 40% identity to known AMPs to minimize homology bias, 56,214 unique peptides remained. Screening this dataset with MultiAMP identified 2,341 putative AMPs, including 484 high-confidence candidates with prediction scores above 0.9 (0.9% of the total screened set).

**Figure 4:**
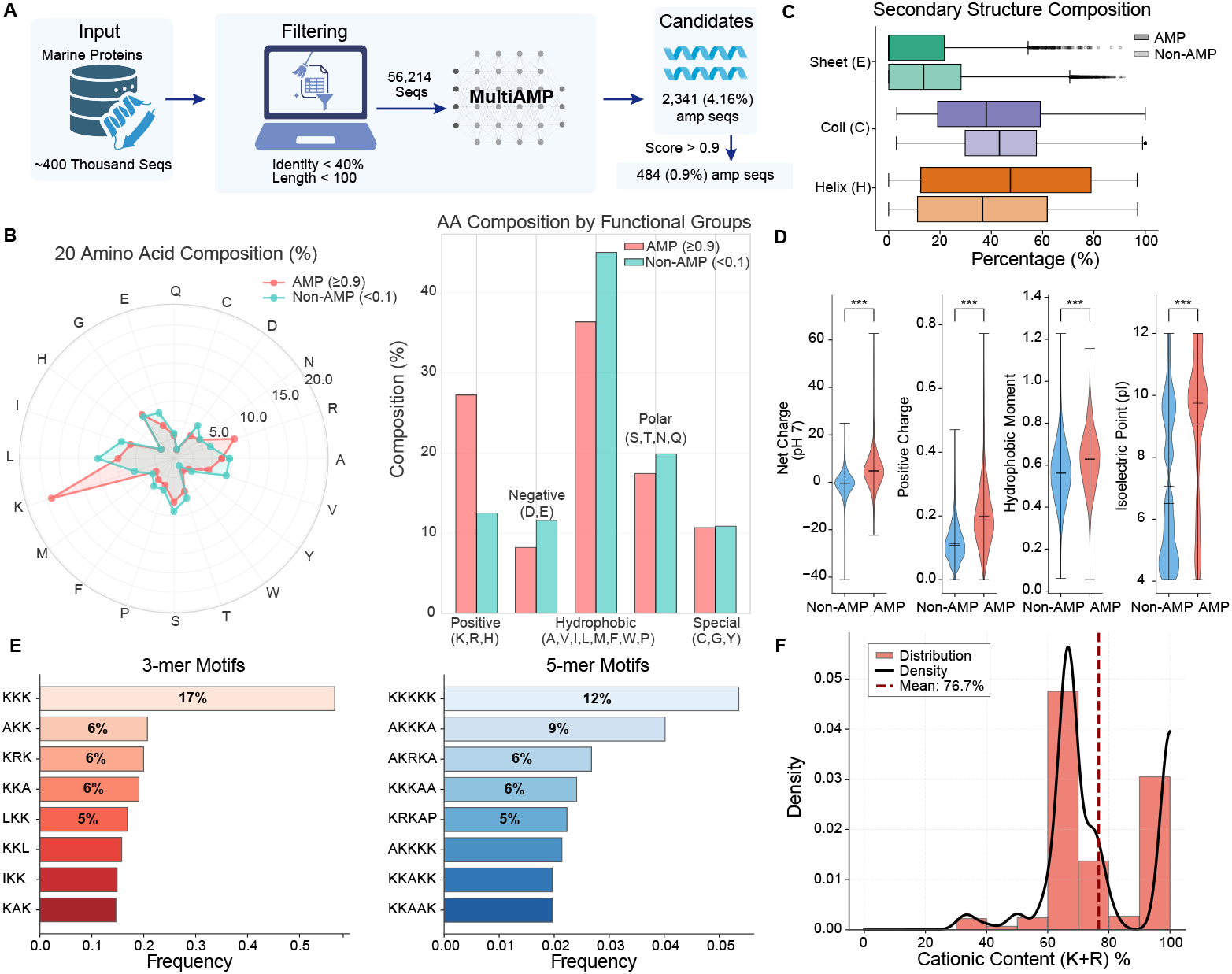
Ocean life mining reveals compositionally distinct AMPs. **(A)** Discovery pipeline: 56,214 ocean peptides (<100 AA, <40% identity to known AMPs) screened with MultiAMP, yielding 484 high-confidence candidates (prediction score > 0.9). **(B)** Amino acid composition comparison (radar plot and grouped bars) of candidate AMPs, which show marked enrichment in positive charge (K+R: 27.2% vs 12.1%) and reduced negative charge (D+E: 7.8% vs 19.3%). **(C)** Secondary structure distributions of candidate AMPs. They exhibit elevated helix (46.0% vs 37.9%), with reduced sheet content (12.2% vs 17.0%) and coil regions compared to predicted non-AMPs. **(D)** Physicochemical properties. AMPs show significantly higher net charge (+5.8 vs +0.3), positive charge ratio (0.27 vs 0.12), hydrophobic moment (0.62 vs 0.48), and isoelectric point (10.2 vs 6.8). **(E)** Enriched motifs discovered from candidate AMPs. Top 3-mer: KKK (17%); top 5-mer: KKKKK (12%). 98.3% of enriched motifs are cationic-rich (≥2 K/R residues). **(F)** Cationic content distribution. AMPs peak at 76.7% K+R content versus 40-50% in non-AMPs.

Analysis of amino acid composition revealed a pronounced compositional bias distinguishing predicted AMPs from non-AMP sequences (Figure 4B). AMP candidates were strongly enriched in positively charged residues, particularly lysine (K) and arginine (R) (combined 27.2% versus 12.1% in non-AMPs), while acidic residues, aspartic acid (D) and glutamic acid (E), were comparatively depleted. This shift produced an overall cationic profile typical of membrane-active AMPs. In addition, hydrophobic residues (F, L, I, M, W) were moderately enriched, contributing to an elevated amphiphilic balance. Predicted structural propensities indicated that ocean-derived AMPs are enriched in *α*-helical content but exhibit reduced *β*-sheet composition relative to non-AMPs (Figure 4C).

Analysis of the physicochemical properties found that they are consistent with sequence-level enrichment (Figure 4D). AMP candidates exhibited a markedly higher net positive charge, positive charge ratio, and isoelectric point, reinforcing their classification as cationic, amphipathic peptides. Motif analysis further highlighted recurrent cationic sequence motifs, with KKK and AKK among the most frequent 3-mers, and KKKKK dominating the 5-mer set (Figure 4E). Nearly all enriched motifs (98.3%) contained at least two consecutive basic residues, reflecting a strong selection for charge clustering. Through the statistics of cationic content distribution (K+R fraction), we confirmed a bimodal bias distinguishing AMPs from non-AMPs, with AMPs exhibiting a mean K+R content of 76.7% compared to 40% in non-AMP sequences (Figure 4F).

Together, these results demonstrate that ocean biological data mining uncovers a sequence distinct class of candidate AMPs, enriched for cationic motifs, amphipathic potential, and membrane-active structural features, suggesting rich untapped diversity within marine peptide space.

### 3.5 Rational design of antimicrobial peptides with specified functional properties

In addition to prediction, MultiAMP enables gradient-based optimization of peptide sequences to achieve desired antimicrobial properties while satisfying biochemical constraints. We implemented three design strategies, *de novo*, motif-guided, and structure-guided to demonstrate the capability for controllable peptide generation (Figure 5A). We first design 10 AMPs each using the *de novo* and motif-guided design strategies (Detailed protocol can be found in Supplementary Note **3**). We evaluated the resulting sequences by using APEX [9] as the oracle, a model trained and validated on extensive in-house AMP datasets to ensure robust generalization to clinically relevant ESKAPE pathogens. The generated peptides exhibited marked MIC reductions across diverse bacterial strains compared with their initial sampled sequences (Figure 5B and C), with up to 7.7-fold improvements and consistent activity gains.

**Figure 5:**
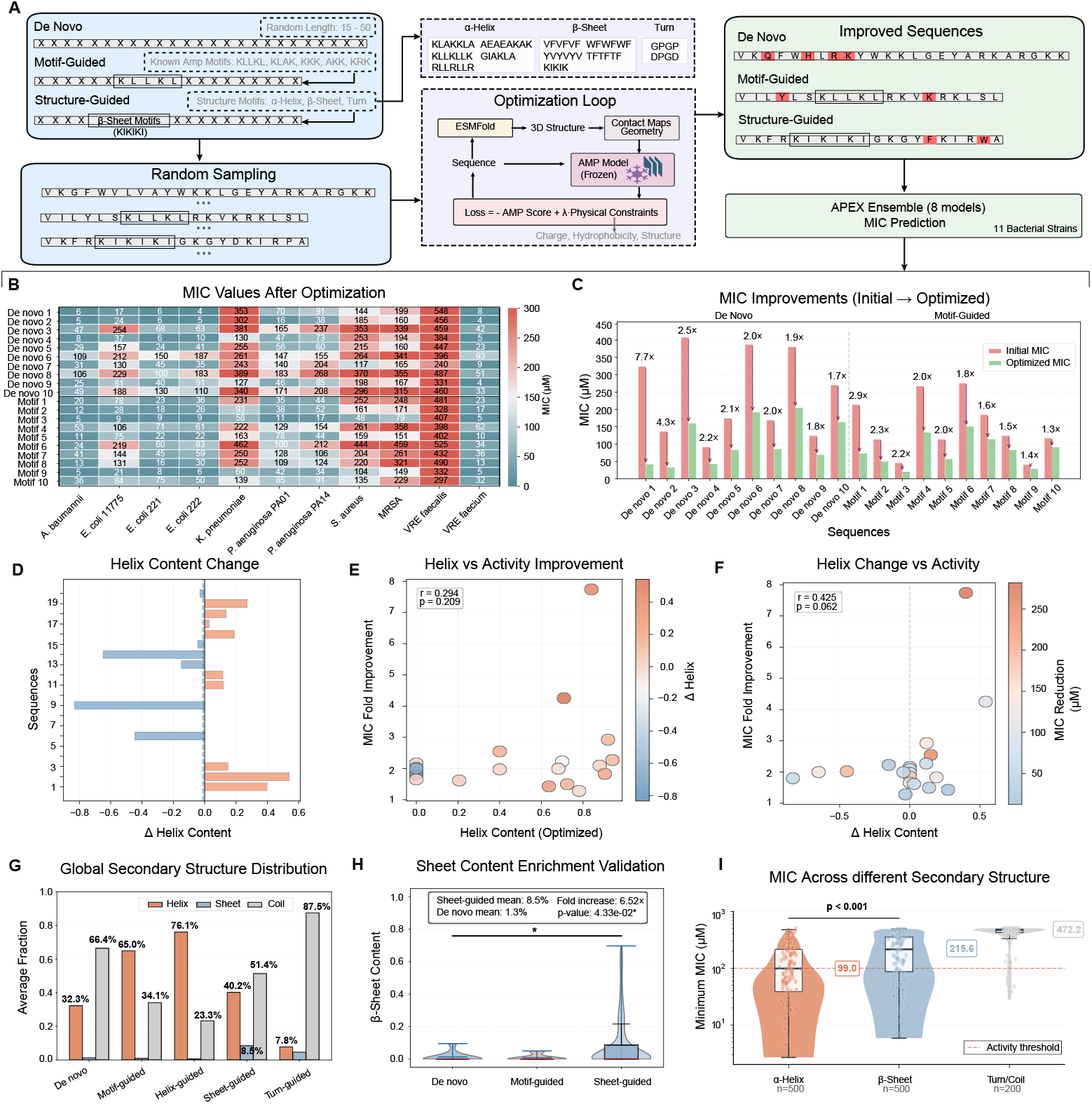
Conditional design enables targeted AMP optimization with enhanced activity. **(A)** We employ gradient-based optimization of peptide sequences for AMP design with MultiAMP. It enables three design strategies: *de novo*, motif-guided, and structure-guided. **(B)** MIC values heatmap for 10 *de novo* and 10 motif-guided designed AMPs. After optimization, the peptides show broad-spectrum activity improvements. **(C)** Fold increase in antimicrobial potency (based on MIC values) after optimization. **(D)** For 20 designed AMPs, we count their helix content changes. **(E)** AMPs with higher helix residues show moderate association with MIC fold improvement. **(F)** An increase in helix content correlates with enhanced antimicrobial potency, reflected by greater MIC reduction. **(G)** Global secondary structure distribution shows that structure-guided designs achieve targeted structure types enrichment. **(H)** *β*-sheet enrichment validation: structure-guided *β*-sheet designs show significantly elevated sheet content (mean 43%) versus *de novo* and motif-guided designs (23-34%). **(I)** MIC values across structural archetypes: *α*-helix and *β*-sheet designs achieve lower MIC. The reported MIC values for each datapoint are the smallest among 11 bacterial strains.

We also counted the Helix residues changes of 20 peptides after optimization (Figure 5D). Peptides with higher optimized helix content tended to display stronger antimicrobial enhancement (Figure 5E), while the extent of helix content change (Δ helix) correlated positively with MIC reduction (Figure 5F). These observations suggest that increased helicity contributes to improved peptide–membrane interactions, and MultiAMP could generate improved AMPs.

Then, for structure-guided design, we generated 500, 500, and 200 sequences with *α*-helix, *β*-sheet, and turn/coil structural fragment constraints, respectively. Representative tertiary structures of designed peptides for each structural category are shown in Supplementary Figure **2**. The secondary-structure profiling of generated peptides confirms the design controllability of MultiAMP: structure-guided optimization produced peptides enriched in target conformations, as shown in Figure 5G and H. Through MIC prediction for these generated AMPs, we found *α*-helical and *β*-sheet peptides generally achieved lower MIC values than those with turn or coil structures (Figure 5I), indicating that explicit structural guidance can effectively bias design outcomes toward more active conformational states. MultiAMP enables the condition-aware, mechanism-informed design of potent and structurally tailored AMPs.

## 4 Discussion

We present MultiAMP, an integrative multi-scale sequence and structure learning framework for AMP prediction and design. By combining multi-level sequence embeddings, geometric structural representations, and secondary-structure-aware regularization, MultiAMP achieves robust generalization on both in-domain and out-of-domain data. Beyond prediction accuracy, the framework offers biologically interpretable insights into AMP structure–function relationships and facilitates rational peptide design.

Unlike traditional models relying solely on amino acid composition or sequence embeddings, Multi-AMP jointly models sequence and structure to capture biologically relevant motifs, serving as both a predictive and hypothesis-generating tool for functional motif discovery and peptide engineering. Application to large-scale ocean-derived data demonstrates its efficiency and ability to uncover previously uncharacterized AMPs with distinct sequence signatures and conserved physicochemical features, enriching our understanding of natural antimicrobial diversity.

While MultiAMP has achieved state-of-the-art performance, future work could extend it to pathogen-specific prediction, MIC estimation, and conditionally-aware modeling of peptide-microbe interactions. Broader validation across organisms and resistance profiles will further establish its generalizability and practical utility. Overall, MultiAMP serves as a powerful, interpretable, and structure-aware framework for AMP discovery and design.

## 5 Acknowledgements

This study was supported by the Shenzhen Medical Research Fund (A2503002 to Y.L.), the National Key R&D Program of China (Grant No.2025YFA0923500 to Y.L.), the Chinese University of Hong Kong (CUHK; award numbers 4937025, 4937026, 5501517, 5501329 and SHIAE BME-p1-24 to Y.L.), the IdeaBooster Fund (IDBF23ENG05 and IDBF24ENG06 to Y.L.), a grant from the Research Grants Council of the Hong Kong Special Administrative Region (HKSAR), China (project numbers CUHK 24204023 and 14208525 to Y.L.), and grants from the Innovation and Technology Commission of the HKSAR, China (project numbers GHP/065/21SZ, ITS/247/23FP and PRP/033/24FX to Y.L.). This research was also partially supported by the Research Matching Grant Scheme at CUHK (award numbers 8601603 and 8601663 to Y.L.) from the Research Grants Council, HKSAR, China.

## Notes

https://github.com/jiayili11/multi-amp

### Competing Interest Statement

The authors have declared no competing interest.

### Summary of Updates

The manuscript has been updated to add a complete Acknowledgments section and to provide detailed funding information, including the relevant funding bodies and award or project numbers.

