## supplementary materials for "Integrative Multi-Scale Sequence–Structure Modeling for Antimicrobial Peptide Prediction and Design"

#### List of Supplementary Notes

**1 Supplementary Note 1: Model Architecture and Training Details** **3**

### List of Supplementary Tables

1 **Comprehensive performance comparison of MultiAMP and baseline methods across all data and similarity-stratified subsets.** . . . 8
2 **Ablation study: Performance of MultiAMP variants with removed components across all data and similarity-stratified subsets.** . . . 9

### 1 Supplementary Note 1: Model Architecture and Training Details

This section provides comprehensive technical details of the MultiAMP architecture, loss function formulations, and training configurations referenced in the main text.

#### 1.1 Architecture Specifications

##### 1.1.1 ESM-2 Multi-layer Aggregation

To obtain multi-scale evolutionary information, we extract and aggregate amino acid representations from specific layers of the ESM-2 (650M) model using learnable layer-specific weights:

$$\mathbf{H}_{esm} = \sum_{l \in \mathcal{L}} w_l \cdot \text{ESM-2}_l(\mathbf{X}), \quad (1)$$

where $\mathbf{X}$ represents the tokenized protein sequence, $\mathcal{L} = \{6, 12, 18, 24, 33\}$ are the selected layer indices, and $w_l$ are learnable weights normalized via softmax such that $\sum_{l \in \mathcal{L}} w_l = 1$. These weights are jointly optimized during training to dynamically emphasize the most informative layer representations for AMP classification. Early layers (6, 12) capture local patterns and amino acid properties, middle layers (18, 24) encode secondary structure and domain-level features, and deep layers (33) capture high-level semantic and functional information.

##### 1.1.2 BiLSTM Sequential Encoder

A bidirectional LSTM captures explicit sequential dependencies:

$$\mathbf{H}_{raw} = [\overrightarrow{LSTM}(\mathbf{X}); \overleftarrow{\text{LSTM}}(\mathbf{X})] \in \mathbb{R}^{L \times 2d_{lstm}}, \quad (2)$$

where $L$ is the sequence length and $d_{lstm} = 128$ is the hidden size per direction, yielding 256-dimensional contextualized embeddings per position.

##### 1.1.3 GVP-GNN Geometric Encoder

For structural encoding, the GVP-GNN processes both scalar and vector geometric features derived from ESMFold-predicted 3D structures. For each residue $i$:

**Initial scalar features** $\mathbf{s}_i^{(0)} \in \mathbb{R}^{d_s}$ include:

- One-hot amino acid identity (20-dim);
- Backbone dihedral angles: $\phi, \psi, \omega$ (3-dim, sine/cosine encoded as 6-dim);
- Sequence position embedding (32-dim sinusoidal encoding);
- Total: $d_s = 58$ dimensions.

**Initial vector features** $\mathbf{v}_i^{(0)} \in \mathbb{R}^{d_v \times 3}$ encode geometric orientations:

- $C_\alpha \rightarrow C_\beta$ unit vector (backbone direction);
- $C_\alpha$(i) $\rightarrow$ $C_\alpha$(i+1) unit vector (chain direction);
- Backbone normal vector: cross product of consecutive $C_\alpha$ directions;
- Local tangent vector: normalized sum of incoming and outgoing $C_\alpha$ directions;
- Total: $d_v = 4$ vector features per residue.

Graph edges connect residues within a spatial distance threshold of 10Å (measured between $C_\alpha$ atoms), with edge features encoding relative distances and orientations.

The GVP-GNN module consists of 3 stacked layers with hidden dimensions $d_s = 384$ (scalar) and $d_v = 4$ (vector). For each GVP-GNN layer, the rotation-equivariant feature update is:

$$\mathbf{s}_{out} = \text{LayerNorm}(\mathbf{W}_{ss}\mathbf{s} + \mathbf{W}_{vs}||\mathbf{v}||_2), \tag{3}$$

$$\mathbf{v}_{out} = \mathbf{W}_{vv}\mathbf{v} \odot \sigma(\mathbf{W}_{sv}\mathbf{s}), \tag{4}$$

where $\mathbf{s} \in \mathbb{R}^{d_s}$ and $\mathbf{v} \in \mathbb{R}^{d_v \times 3}$ denote the scalar and vector features; $\mathbf{W}_{ss} \in \mathbb{R}^{d_s \times d_s}$, $\mathbf{W}_{vs} \in \mathbb{R}^{d_s \times d_v}$, $\mathbf{W}_{vv} \in \mathbb{R}^{d_v \times d_v}$, and $\mathbf{W}_{sv} \in \mathbb{R}^{d_v \times d_s}$ are learnable weight matrices; $|\mathbf{v}|_2 \in \mathbb{R}^{d_v}$ computes the L2 norm of each vector feature; $\sigma(\cdot)$ is the SiLU activation; and $\odot$ represents element-wise multiplication. This architecture maintains rotation equivariance: if the input coordinates are rotated by $\mathbf{R} \in SO(3)$, the output vector features transform as $\mathbf{v}_{out} \to \mathbf{R}\mathbf{v}_{out}$ while scalar features remain invariant.

##### 1.1.4 Cross-attention Fusion

Multi-head cross-attention (8 heads, dimension 512) aligns ESM-2 and BiLSTM features:

$$\mathbf{H}_{cross} = \text{MultiHeadAttn}(\mathbf{H}_{esm}, \mathbf{H}_{raw}, \mathbf{H}_{raw}) + \mathbf{H}_{esm}, \tag{5}$$

where $\mathbf{H}_{esm}$ serves as queries (evolutionary context) and $\mathbf{H}_{raw}$ as keys and values (local sequential patterns), followed by a residual connection.

##### 1.1.5 Deep Fusion Transformer

A 3-layer transformer encoder (8 heads, hidden dimension 512, feedforward dimension 2048) integrates cross-attended sequence features and GVP-GNN structural embeddings:

$$\mathbf{H}_{fused} = \text{TransformerEncoder}([\mathbf{h}_{cls}; \mathbf{H}_{cross} + \mathbf{H}_{struct}]), \tag{6}$$

where $\mathbf{h}_{cls}$ is the CLS token from ESM-2, and $\mathbf{H}_{struct}$ are node embeddings from GVP-GNN's final layer.

##### 1.1.6 Gated Fusion Mechanism

Final representation combines sequence-level (CLS token from transformer output, $\mathbf{h}_{cls}^{final} \in \mathbb{R}^{512}$) and graph-level features (global mean pooling from GVP-GNN, $\mathbf{g}_{global} \in \mathbb{R}^{384}$) via learned gating:

$$\mathbf{f}_{final} = [\mathbf{h}_{cls}^{final}; \mathbf{g}_{global}] \in \mathbb{R}^{896}, \tag{7}$$

$$\mathbf{z} = \mathbf{f}_{final} \odot \sigma(\mathbf{W}_g \mathbf{f}_{final}), \tag{8}$$

where $\mathbf{W}_g \in \mathbb{R}^{896 \times 896}$ is a learnable gating matrix, $\sigma(\cdot)$ is sigmoid activation, and $\odot$ is element-wise multiplication. The gate learns to adaptively weight sequence vs. structure contributions based on input characteristics.

#### 1.2 Loss Function Formulations

##### 1.2.1 Primary Classification Loss

The model employs standard Binary Cross-Entropy (BCE) loss for AMP binary classification:

$$\mathcal{L}_{cls} = \text{BCE}(\sigma(\mathbf{W}_c \mathbf{z}), y), \tag{9}$$

where $\mathbf{z} \in \mathbb{R}^d$ represents the finalized embedding extracted by our network for a given input sample, $\mathbf{W}_c \in \mathbb{R}^{1 \times d}$ is the classification weight matrix, $\sigma(\cdot)$ is the sigmoid activation function producing predicted probability $\hat{y} = \sigma(\mathbf{W}_c \mathbf{z})$, and $y \in \{0, 1\}$ is the ground-truth binary label (1 for AMP, 0 for non-AMP).

###### 1.2.2 Supervised Contrastive Loss

To sculpt a more structured and discriminative embedding space, we incorporate supervised contrastive learning. This objective encourages the model to pull embeddings of samples from the same class (positives) closer together while pushing different-class samples apart. The loss for a given anchor sample $\mathbf{z}_i$ is defined as:

$$\mathcal{L}_{contrast} = -\log \frac{\exp(\text{sim}(\mathbf{z}_i, \mathbf{z}_j^+)/\tau)}{\sum_{k\in\mathcal{B}\setminus\{i\}} \exp(\text{sim}(\mathbf{z}_i, \mathbf{z}_k)/\tau)}, \tag{10}$$

where:

- $\text{sim}(\mathbf{u}, \mathbf{v}) = \frac{\mathbf{u}^T\mathbf{v}}{||\mathbf{u}||_2 \cdot ||\mathbf{v}||_2}$ denotes cosine similarity;
- $\mathbf{z}_j^+$ represents a randomly sampled positive sample with the same label as the anchor within the batch;
- $\mathcal{B}$ denotes the set of all samples in the current training batch (including both AMPs and non-AMPs);
- $k$ indexes all samples in $\mathcal{B}$ except the anchor itself;
- $\tau = 0.07$ is the temperature parameter controlling the concentration of the distribution.

Each training batch is constructed by randomly sampling peptides with balanced class distribution (equal numbers of AMPs and non-AMPs, batch size 16 with 8 AMPs and 8 non-AMPs) to ensure sufficient positive pairs for effective contrastive learning. For each anchor, we sample one positive from the same class within the batch as $\mathbf{z}_j^+$, while all other samples in the batch (both same-class and different-class) contribute to the denominator.

###### 1.2.3 Secondary Structure Reconstruction Loss

To infuse the model with structural awareness, we introduce a secondary structure prediction auxiliary task guided by a composite loss function:

$$\mathcal{L}_{ss} = \mathcal{L}_{focal} + \lambda_{crf}\mathcal{L}_{crf} + \lambda_{cont}\mathcal{L}_{continuity}, \tag{11}$$

$$\mathcal{L}_{focal} = -\frac{1}{L}\sum_{i=1}^{L} \alpha_t(1 - p_t)^\gamma \log(p_t), \tag{12}$$

$$\mathcal{L}_{crf} = -\log P(\mathbf{y}_{ss}|\mathbf{X}, \mathbf{mask}), \tag{13}$$

where $\lambda_{crf} = 0.3$ and $\lambda_{cont} = 0.1$ are loss weighting coefficients determined via validation set tuning.

**Focal loss component** addresses severe class imbalance in secondary structure annotations. In typical peptide datasets, coil regions ('C') constitute 60-70% of residues, while helix ('H') and sheet ('E') states are minority classes. The focal loss re-weights samples based on prediction difficulty:

- $p_t$: Model's predicted probability for the true class at position $t$. Specifically, $p_t = p$ if the true class is positive (e.g., helix when ground truth is helix), and $p_t = 1 - p$ otherwise, where $p$ is the raw prediction probability;
- $(1 - p_t)^\gamma$: Modulating factor that down-weights loss contribution from well-classified examples ($p_t \to 1$) and focuses learning on hard examples ($p_t \to 0$). We set $\gamma = 3.0$ based on validation performance;

- $\alpha_t$: Class-balancing weight computed as inverse class frequency normalized across the training set: $\alpha_c = \frac{1/n_c}{\sum_{c'} 1/n_{c'}}$ where $n_c$ is the number of residues with class $c$. Typical values: $\alpha_{coil} = 0.25$, $\alpha_{helix} = 0.75$, $\alpha_{sheet} = 0.75$.

**CRF loss component** captures dependencies between adjacent structural states by modeling the entire sequence jointly rather than position-independently. The CRF learns a transition matrix $\mathbf{T} \in \mathbb{R}^{3\times3}$ where $T_{ij}$ represents the log-probability of transitioning from state $i$ to state $j$ (for H/E/C states). The CRF loss maximizes the conditional log-likelihood:

$$\begin{aligned}\mathcal{L}_{crf} &= -\log P(\mathbf{y}_{ss}|\mathbf{X}, \mathbf{mask}) \\ &= -\left(\sum_{i=1}^{L} \psi_i(y_i) + \sum_{i=1}^{L-1} T_{y_i,y_{i+1}} - \log Z\right),\end{aligned} \tag{14}$$

where $\psi_i(y_i)$ are unary potentials (logits from the model), $T_{y_i,y_{i+1}}$ are learned pairwise transition scores, and $Z$ is the partition function computed via the forward algorithm. The mask handles variable-length sequences by excluding padding positions.

**Continuity regularization** promotes smooth structural predictions and penalizes fragmented, abrupt transitions:

$$\mathcal{L}_{continuity} = \frac{1}{L-1} \sum_{i=1}^{L-1} ||\mathbf{p}_i^{ss} - \mathbf{p}_{i+1}^{ss}||_2^2, \tag{15}$$

where $\mathbf{p}_i^{ss} \in \mathbb{R}^3$ denotes the predicted probability distribution (after softmax) over the three secondary structure states at position $i$, and $||\cdot||_2$ is the L2 norm. This term encourages adjacent positions to have similar structural predictions, reducing unrealistic isolated helix/sheet residues surrounded by coils.

##### 1.2.4 Total Multi-task Objective

The complete training objective combines all components with dynamic weighting:

$$\mathcal{L}_{total} = w_{cls}\mathcal{L}_{cls} + w_{contrast}\mathcal{L}_{contrast} + w_{ss}\mathcal{L}_{ss}, \tag{16}$$

where:

- $w_{cls} = 1.0$ (fixed) for the primary classification task;
- $w_{contrast} = 0.1$ (initial) for contrastive learning;
- $w_{ss} = 0.15$ (initial) for secondary structure prediction.

These auxiliary task weights were selected through grid search on the validation set, exploring values in the ranges $w_{contrast} \in [0.05, 0.2]$ and $w_{ss} \in [0.1, 0.3]$, with the final values yielding the highest validation MCC. To gradually emphasize the primary classification objective while maintaining multi-task regularization benefits throughout training, $w_{contrast}$ and $w_{ss}$ decay by a factor of 0.90 per epoch starting from epoch 25, reaching approximately $w_{contrast} \approx 0.06$ and $w_{ss} \approx 0.09$ by epoch 30.

#### 1.3 Training Configuration

##### 1.3.1 Optimization Strategy

The model was trained using the AdamW optimizer. A differential learning rate was applied, with the pre-trained language model backbone set to $1 \times 10^{-5}$ and the randomly initialized head layers set to $5 \times 10^{-5}$. We employed a Cosine Annealing scheduler with Warm Restarts ($T_0 = 25$, $T_{mult} = 2.0$) and a minimum learning rate of $1 \times 10^{-7}$, preceded by a 5-epoch linear warmup phase. Training was conducted with a batch size of 16 for a maximum of 30 epochs, utilizing an early stopping mechanism based on validation MCC with a patience of 8 epochs. Gradient clipping with a maximum norm of 1.0 was applied to ensure training stability.

##### 1.3.2 Regularization

To mitigate overfitting, we incorporated the following regularization techniques:

- **Dropout**: A rate of 0.3 was applied to all fully connected and attention layers;
- **Weight Decay**: An L2 weight decay (AdamW's decoupled weight decay) of $\lambda = 0.01$ was applied to all trainable parameters.

##### 1.3.3 Data Augmentation

Online data augmentation was applied during training to improve model generalization:

- **Sequence augmentation**: Random amino acid substitution was performed with a 10% probability per residue, using BLOSUM62-weighted sampling to maintain evolutionary and physicochemical plausibility;
- **Structure augmentation**: To enhance structural robustn-ess, random 3D rotations (uniformly sampled from SO(3)) and Gaussian noise ($\sigma = 0.1$ Å) were applied to atomic coordinates.

#### 1.4 Hyperparameter Search

An extensive hyperparameter search was conducted via grid search using 3-fold cross-validation on the training set to determine the optimal model architecture. The search space included key parameters for each module. The final selected configuration, validated on a held-out set, is as follows:

- **GVP-GNN**: 3 layers with 384 hidden scalar dimensions and 4 vector dimensions per node;
- **Bi-LSTM**: 2 layers with 128 hidden units per direction;
- **Transformer Fusion Encoder**: 3 encoder layers with 8 attention heads;
- **ESM-2 Strategy**: The top 6 layers of the pre-trained `esm2_t33_650M_UR50D` model were unfrozen and fine-tuned, while the lower layers remained frozen.

#### 1.5 Computational Requirements

##### 1.5.1 Hardware and Runtime

All experiments were conducted on a workstation equipped with NVIDIA RTX A6000 GPUs (48 GB memory), running on CUDA 12.7 (Driver Version: 565.57.01). Models were implemented in PyTorch and trained using automatic mixed precision (AMP) to accelerate computation. The full model training for 30 epochs took approximately 6 hours. Inference on the test set of 5,355 sequences was highly efficient, completing in under 2 minutes (~45 sequences/second).

##### 1.5.2 Memory Optimization

To manage the memory demands of large protein language models and 3D structures, several optimization strategies were employed:

- **Gradient Checkpointing**: Applied to the ESM-2 model during feature extraction to trade computation for a significant reduction in memory usage;
- **Dynamic Batching**: The batch size was adaptively adjusted based on sequence length to maintain a consistent memory footprint;
- **CPU Offloading**: Pre-computation of structural features, such as those from ESMFold, was performed offline and cached to disk to free up GPU memory for training.

### 2 Supplementary Note 2: Performance Evaluation

**Supplementary Table 1: Comprehensive performance comparison of MultiAMP and baseline methods across all data and similarity-stratified subsets.**

| Subset | Method | AUROC | MCC | Accuracy | F1 | Precision | Recall |
|---|---|---|---|---|---|---|---|
| *All Data (N=5,355)* | | | | | | | |
| | MultiAMP | **0.9810** | **0.8519** | **0.9477** | **0.8857** | **0.8923** | 0.8793 |
| | PepNet | 0.9642 | 0.7640 | 0.9079 | 0.8189 | 0.7488 | **0.9036** |
| | AMPScanner | 0.9553 | 0.7700 | 0.9143 | 0.8243 | 0.7810 | 0.8728 |
| | AMAP | 0.9415 | 0.6983 | 0.8838 | 0.7703 | 0.7076 | 0.8452 |
| | AMP-BERT | 0.8832 | 0.6885 | 0.8838 | 0.7630 | 0.7201 | 0.8112 |
| | Macrel | 0.9235 | 0.6376 | 0.8577 | 0.7251 | 0.6534 | 0.8144 |
| | iAMP-2L | 0.9063 | 0.6278 | 0.8415 | 0.7154 | 0.6101 | 0.8647 |
| | AmPEP | 0.9022 | 0.5881 | 0.8495 | 0.6861 | 0.6604 | 0.7139 |
| *Low Similarity (<40% identity, N=4,379)* | | | | | | | |
| | MultiAMP | **0.9629** | **0.7169** | **0.9491** | **0.7451** | **0.7443** | 0.7460 |
| | PepNet | 0.9199 | 0.5775 | 0.9023 | 0.6137 | 0.5067 | **0.7780** |
| | AMPScanner | 0.8992 | 0.5673 | 0.9103 | 0.6105 | 0.5385 | 0.7048 |
| | AMAP | 0.8977 | 0.4923 | 0.8813 | 0.5398 | 0.4401 | 0.6979 |
| | Macrel | 0.8696 | 0.4208 | 0.8529 | 0.4747 | 0.3688 | 0.6659 |
| | iAMP-2L | 0.8401 | 0.4018 | 0.8299 | 0.4518 | 0.3330 | 0.7025 |
| | AmPEP | 0.8335 | 0.3251 | 0.8525 | 0.3985 | 0.3359 | 0.4897 |
| | AMP-BERT | 0.7785 | 0.4507 | 0.8826 | 0.5077 | 0.4366 | 0.6064 |
| *High Similarity (>=40% identity, N=976)* | | | | | | | |
| | MultiAMP | 0.9799 | **0.8143** | **0.9416** | **0.9638** | **0.9756** | 0.9523 |
| | PepNet | **0.9816** | 0.7692 | 0.9334 | 0.9598 | 0.9474 | **0.9724** |
| | AMPScanner | 0.9741 | 0.7697 | 0.9324 | 0.9589 | 0.9529 | 0.9649 |
| | Macrel | 0.9347 | 0.6415 | 0.8791 | 0.9237 | 0.9533 | 0.8959 |
| | AMAP | 0.9339 | 0.6637 | 0.8955 | 0.9354 | 0.9449 | 0.9260 |
| | AmPEP | 0.9235 | 0.5733 | 0.8361 | 0.8929 | 0.9570 | 0.8369 |
| | AMP-BERT | 0.9119 | 0.6421 | 0.8893 | 0.9316 | 0.9400 | 0.9235 |
| | iAMP-2L | 0.8645 | 0.6223 | 0.8934 | 0.9360 | 0.9190 | 0.9536 |

All metrics were evaluated on the test set (5,355 peptides total) as shown in Supplementary Table 1. **Bold values** indicate the best performance in each subset. F1, Precision, and Recall refer to the positive class (AMP). Low-similarity subset (<40% sequence identity to training data) tests the generalization capability. High-similarity subset (>=40% identity) represents easier classification with closer training analogs. MultiAMP achieves superior performance on all data and low-similarity subsets, demonstrating robust generalization. On high-similarity data, MultiAMP achieves the highest MCC despite slightly lower AUROC than PepNet, indicating better balanced performance across precision and recall.

Key findings found in Supplementary Table 2: (1) On low-similarity data, the Full Model achieves the highest AUROC (0.9629) and MCC (0.7169), with performance degrading most when removing ESM-2 ($\Delta$AUROC = -7.15%) or GVP-GNN ($\Delta$AUROC = -3.26%), demonstrating that both deep sequence features and explicit graph-based structural information are critical for generalization to distant homologs. (2) On high-similarity data, removing GVP-GNN (w/o GVP) surprisingly improves some metrics, suggesting that for peptides with close training analogs, deep sequence features dominate and structural information may introduce minor noise. (3) Removing ESM-2 causes the most severe degradation across all subsets (AUROC drops by 3.9-7.2%), confirming its role as the primary feature extractor. (4) Bi-LSTM provides complementary shallow sequential pattern learning: removing it causes modest performance drops on low-similarity data ($\Delta$AUROC = -0.08%, $\Delta$MCC = -0.02) but minimal impact on high-similarity data, indicating it aids in capturing local motifs when training analogs are distant. (5) The secondary structure prediction task functions as a crucial geometric regularizer. While removing

**Supplementary Table 2: Ablation study: Performance of MultiAMP variants with removed components across all data and similarity-stratified subsets.**

| Subset | Model | AUROC | MCC | Accuracy | F1 | Precision | Recall |
|---|---|---|---|---|---|---|---|
| *All Data (N=5,355)* | | | | | | | |
| | Full Model | **0.9810** | **0.8519** | **0.9477** | **0.8857** | 0.8923 | **0.8793** |
| | w/o SS | 0.9747 | 0.8515 | 0.9425 | 0.8830 | **0.9262** | 0.8436 |
| | w/o BiLSTM | 0.9794 | 0.8396 | 0.9430 | 0.8766 | 0.8755 | 0.8776 |
| | w/o GVP+SS | 0.9739 | 0.8287 | 0.9318 | 0.8684 | 0.8712 | 0.8657 |
| | w/o GVP | 0.9702 | 0.8367 | 0.9396 | 0.8703 | 0.8745 | 0.8662 |
| | w/o GVP+SS+BiLSTM | 0.9680 | 0.8293 | 0.9373 | 0.8627 | 0.8698 | 0.8558 |
| | w/o ESM2 | 0.9420 | 0.7406 | 0.9107 | 0.7957 | 0.8418 | 0.7545 |
| *Low Similarity (<40% identity, N=4,379)* | | | | | | | |
| | Full Model | **0.9629** | **0.7169** | **0.9491** | **0.7451** | 0.7443 | 0.7460 |
| | w/o SS | 0.9514 | 0.7168 | 0.9420 | 0.7401 | **0.8059** | 0.6842 |
| | w/o BiLSTM | 0.9621 | 0.6968 | 0.9436 | 0.7277 | 0.7021 | **0.7551** |
| | w/o GVP+SS | 0.9443 | 0.6854 | 0.9398 | 0.7130 | 0.6935 | 0.7337 |
| | w/o GVP | 0.9303 | 0.6498 | 0.9331 | 0.6841 | 0.6689 | 0.6999 |
| | w/o GVP+SS+BiLSTM | 0.9307 | 0.6400 | 0.9308 | 0.6768 | 0.6612 | 0.6931 |
| | w/o ESM2 | 0.8914 | 0.5397 | 0.9208 | 0.5824 | 0.6142 | 0.5538 |
| *High Similarity (>=40% identity, N=976)* | | | | | | | |
| | Full Model | 0.9799 | 0.8143 | 0.9416 | 0.9638 | 0.9756 | 0.9523 |
| | w/o SS | 0.9737 | 0.8015 | 0.9324 | 0.9574 | **0.9854** | 0.9310 |
| | w/o BiLSTM | 0.9810 | 0.8175 | 0.9406 | 0.9579 | 0.9598 | 0.9560 |
| | w/o GVP+SS | 0.9810 | **0.8333** | 0.9488 | 0.9630 | 0.9612 | 0.9648 |
| | w/o GVP | **0.9850** | 0.8306 | **0.9508** | **0.9647** | 0.9624 | **0.9670** |
| | w/o GVP+SS+BiLSTM | 0.9774 | 0.8021 | 0.9365 | 0.9605 | 0.9687 | 0.9525 |
| | w/o ESM2 | 0.9503 | 0.6412 | 0.8658 | 0.9132 | 0.9677 | 0.8645 |

**Model variants:** Full Model = complete MultiAMP architecture; w/o SS = secondary structure prediction task removed; w/o GVP = GVP-GNN structure encoder removed; w/o GVP+SS = GVP-GNN and secondary structure prediction task removed; w/o GVP+SS+ BiLSTM = GVP-GNN, secondary structure prediction, and Bi-LSTM removed (ESM-2 only for sequence); w/o BiLSTM = Bi-LSTM shallow sequence encoder removed; w/o ESM2 = ESM-2 deep sequence encoder removed. Bi-LSTM captures local sequential patterns while ESM-2 provides deep evolutionary context. **Bold values** indicate the best performance in each subset.

this task (w/o SS) yields higher Precision across all subsets (e.g., rising from 0.7443 to 0.8059 on low-similarity data), it causes a significant drop in Recall (falling from 74.60% to 68.42%). This trade-off indicates that without structural constraints, the model adopts a conservative strategy, likely overfitting to explicit sequence motifs; the multi-task objective forces the learning of underlying structure-function dependencies, thereby maintaining high sensitivity for identifying diverse AMPs with low sequence homology.

### 3 Supplementary Note 3: Gradient-Based Sequence Optimization

**Sequence representation** Each peptide sequence of length $L$ is represented as a soft one-hot matrix $\mathbf{S} \in \mathbb{R}^{L\times 20}$, where each row $\mathbf{s}_i$ is a probability distribution over the 20 canonical amino acids. This representation is initialized randomly for de novo design or from template sequences for motif-guided and structure-guided approaches.

**Optimization objective** The optimization maximizes a composite loss function balancing antimicrobial activity prediction, physicochemical constraints, and structural objectives:

$$\mathcal{L} = w_{amp} \cdot \sigma(f_\theta(\mathbf{S})) - w_{phys} \cdot \mathcal{R}_{phys}(\mathbf{S}) - w_{struct} \cdot \mathcal{R}_{struct}(\mathbf{S}), \tag{17}$$

where:

- $f_\theta(\mathbf{S})$: MultiAMP classifier logits (frozen parameters);
- $\sigma(\cdot)$: Sigmoid activation to obtain AMP probability;
- $\mathcal{R}_{phys}(\mathbf{S})$: Physicochemical property regularization (charge, hydrophobicity, amphipathicity);
- $\mathcal{R}_{struct}(\mathbf{S})$: Secondary structure constraint term (structure-guided only);
- Loss weights: $w_{amp} = 1.0$, $w_{phys} = 0.3$, $w_{struct} = 0.5$.

**Physicochemical regularization** The physicochemical regularization term enforces biologically plausible properties:

$$\begin{aligned}\mathcal{R}_{phys}(\mathbf{S}) =& \lambda_{charge} \cdot |\text{Charge}(\mathbf{S}) - \text{Charge}_{target}|^2 + \\ & \lambda_{hydro} \cdot |\text{GRAVY}(\mathbf{S})|^2 + \lambda_{div} \cdot H(\mathbf{S}),\end{aligned} \tag{18}$$

where:

- $\text{Charge}(\mathbf{S})$: Net charge calculated as $\sum_i \mathbf{s}_i^T \mathbf{c}$, with $\mathbf{c}$ being amino acid charge vector;
- $\text{Charge}_{target}$: Target charge range (+3 to +6 for AMPs);
- $\text{GRAVY}(\mathbf{S})$: Grand average of hydropathy index;
- $H(\mathbf{S})$: Shannon entropy encouraging amino acid diversity;
- $\lambda_{charge} = 0.5$, $\lambda_{hydro} = 0.2$, $\lambda_{div} = 0.1$.

**Gumbel-Softmax sampling** To enable discrete sequence generation while maintaining differentiability, we apply Gumbel-Softmax [1] reparameterizat-ion:

$$\mathbf{s}_i = \text{softmax}\left(\frac{\log \mathbf{p}_i + \mathbf{g}}{\tau}\right), \tag{19}$$

where $\mathbf{p}_i$ are learned logits, $\mathbf{g} \sim \text{Gumbel}(0, 1)$, and temperature $\tau$ anneals from 1.0 to 0.1 over 50 iterations.

**Optimization procedure** Sequences are optimized using the Adam optimizer (learning rate: 0.01) for 30-50 iterations:

1. Initialize $\mathbf{S}$ (random for de novo, template-based for motif/structure-guided)
2. For each iteration:
   - Sample discrete sequence via Gumbel-Softmax;
   - Predict 3D structure using ESMFold;
   - Compute MultiAMP score $f_\theta(\mathbf{S})$;
   - Evaluate physicochemical properties and structural constraints;
   - Compute total loss $\mathcal{L}$;
   - Update $\mathbf{S}$ via gradient ascent.
3. Select top sequences by AMP score for downstream validation.

##### 3.1 Strategy-Specific Details

**De novo design**

- **Initialization**: Random amino acid probabilities sampled from a uniform distribution over 20 canonical amino acids, optionally biased toward AMP-favorable composition
- **Initialization strategies (5 approaches to ensure diversity)**:
  - Random short (15-25 residues): Uniform random initialization;
  - Random medium (25-35 residues): Uniform random initialization;
  - Random long (35-45 residues): Uniform random initialization;
  - Cationic-rich: Biased initialization with K (12%), R (8%) enrichment;
  - Hydrophobic-rich: Biased initialization with L (10%), I (8%), V (8%), A (8%) enrichment.
- **Length distribution**: 20-30 residues (varied across designs to span short-to-medium AMP length range).
- **Constraints**: Only physicochemical regularization applied:
  - Charge constraint: Target net charge +3 to +6 (typical for AMPs);
  - Hydrophobicity constraint: GRAVY score penalty to avoid extreme values;
  - Diversity constraint: Shannon entropy term prevents sequence simplification.
- **Design scale**: Generated 500 candidate sequences (100 per initialization strategy), selected the top 10 by AMP score for APEX validation.

**Motif-guided design**

- **Functional motif library**: 5 sequence motifs selected based on Figure 4 enrichment analysis and classical AMP literature:
  - **KKK** (3 residues): Top enriched motif from ocean AMP discovery (Figure 4E), appears in 56.8% of high-scoring sequences;
  - **KRK** (3 residues): Cationic motif from Figure 4 analysis, found in 20.0% of ocean AMPs;
  - **AKK** (3 residues): Lysine-rich motif from Figure 4, present in 20.7% of predicted AMPs;
  - **KLLKL** (5 residues): Classical magainin-like amphipathic motif with hydrophobic leucines alternating with cationic lysines;
  - **KLAK** (4 residues): Canonical amphipathic motif from the cecropin family, widely used in AMP design.
- **Initialization**: Template sequences (20-40 residues) with motif embedded at random positions (5-15 residues from termini to avoid edge effects);
- **Motif preservation**: Hard constraint via binary mask-motif positions receive fixed one-hot encodings during optimization, gradients blocked for these positions;
- **Optimization scope**: Only flanking residues (prefix and suffix) updated via gradient descent; motif region remains unchanged throughout 40 iterations;
- **Flanking sequence bias**: Random initialization with slight enrichment in AMP-favorable residues (K: 12%, R: 8%, L: 10%, A: 8%);
- **Design scale**: Generated 100 variants per motif (500 total sequences), selected the best variant per motif based on AMP score, yielding 5 final candidates for APEX validation.

**Structure-guided design**

- **Structural motif library**: 12 sequence fragments with experimentally validated secondary structure propensities, organized by target structure:
  - **$\alpha$-Helix motifs (5 total, 6-8 residues)**:
    * KLAKKLA (7 aa): Amphipathic helix with alternating cationic (K) and hydrophobic (L, A) residues; magainin-like membrane insertion motif;
    * KLLKLLK (7 aa): Leucine-rich amphipathic helix with strong hydrophobic face (LLL) and cationic face (KK); enhanced membrane affinity;
    * RLLRLLR (7 aa): Arginine-leucine helix; arginine provides stronger electrostatic interactions than lysine for bacterial membrane targeting;
    * AEAEAKAK (8 aa): Charged residue helix balancing anionic (E) and cationic (K) for pH-dependent activity;
    * GIAKLA (6 aa): Short compact helix with glycine flexibility for membrane insertion.
  - **$\beta$-Sheet motifs (5 total, 6 residues)**:
    * VFVFVF (6 aa): Alternating valine-phenylalanine; strong $\beta$-sheet propensity via hydrophobic stacking;
    * YVYVYV (6 aa): Tyrosine-valine aromatic strand; $\pi$-$\pi$ stacking enhances sheet stability;
    * KIKIKI (6 aa): ALysine-isoleucine alternation; amphiphilic $\beta$-sheet with charged edges and a hydrophobic core;
    * WFWFWF (6 aa): Tryptophan-rich strand for membrane anchoring via indole side chains;
    * TFTFTF (6 aa): Threonine-phenylalanine; hydrogen bonding network stabilizes sheet structure.
  - **Turn motifs (2 total, 4 residues)**:
    * GPGP (4 aa): Glycine-proline repeats; proline induces kink, glycine provides flexibility for tight turns;
    * DPGD (4 aa): Charged turn motif with aspartate flanking; promotes $\beta$-turn formation.
- **Initialization**: Structural motif inserted at random positions, 3-15 residues from termini, within 20-40 residue templates; total sequence length varied to prevent length bias;
- **Flanking sequence generation**: Non-motif regions biased toward structure-compatible amino acids using secondary structure propensity scales:
  - $\alpha$-Helix flanking (70% of designs): Enriched in A (15%), E (12%), L (12%), M (8%), K (12%), R (8%)—high helix propensity residues;
  - $\beta$-Sheet flanking (70% of designs): Enriched in V (15%), I (12%), F (10%), Y (8%), W (6%), T (10%)—high sheet propensity residues;
  - Turn flanking (70% of designs): Enriched in G (15%), P (10%), D (8%), S (8%), N (8%)—high turn propensity residues;
  - Random flanking (30% of designs): AMP-like composition without structure bias for diversity.
- **Motif preservation during optimization**: Hard constraint via binary mask—structural motif positions fixed with one-hot encodings throughout 50 iterations; only flanking residues optimized;

- **Design scale**: Generated 100 variants per structural motif × 12 motifs = 1,200 total sequences;
- **Post-optimization validation**: ESMFold 3D structure prediction for all 1,200 sequences, followed by Ramachandran-based secondary structure assignment:
  - $\alpha$**-Helix**: Phi/psi angles in range ($-90 < \phi < -30$, $-70 < \psi < -20$) assigned as H;
  - $\beta$**-Sheet**: Phi/psi angles outside helix range with extended conformation assigned as E;
  - **Coil/Turn**: Remaining angles assigned as C.
- **Success criteria (validated post-hoc on motif region)**:
  - $\alpha$-Helix success: $>70\%$ of motif residues adopt H conformation; 5 motifs tested, success rates 68-89%;
  - $\beta$-Sheet success: $>50\%$ of motif residues adopt E conformation; 5 motifs tested, success rates 52-78%;
  - Turn success: $>60\%$ of motif residues adopt C conformation; 2 motifs tested, success rates 71-85%.
- **Final selection**: From 1,200 total designs, selected the top 20 by AMP score for APEX validation; analyzed all structures for Supplementary Figure **??**A-C validation panels.

#### 3.2 Activity Prediction and Validation

**MIC prediction via APEX ensemble** All optimized sequences were evaluated utilizing the APEX [2] ensemble model to predict minimum inhibitory concentrations (MIC) for 11 bacterial strains:

- **Gram-negative**: *E. coli*, *A. baumannii*, *P. aeruginosa*, *K. pneumoniae*;
- **Gram-positive**: *S. aureus*, *E. faecalis*, *MRSA*;
- **Other pathogens**: *C. albicans*, *E. faecium*, *S. epidermidis*, *S. pneumoniae*.

APEX predictions were averaged across 8 ensemble models to obtain robust MIC estimates. Activity improvement was quantified as fold-change: Fold $=$ MIC$_{initial}$/MIC$_{optimized}$.

**Structural analysis** Post-optimization structural characterization included:

- **ESMFold prediction**: Full 3D structure prediction for all candidates;
- **Secondary structure extraction**: Automated DSSP-like analysis from PDB coordinates;
- **Helix content change**: Quantified as $\Delta H = H_{optimized} - H_{initial}$ (Figure 5D);
- **Structure-activity correlation**: Pearson correlation between structural features and MIC improvement (Figure 5E and F).

**Diversity and novelty assessment** To ensure designed sequences explore novel chemical space:

- **Sequence similarity**: BLAST against known AMP databases (DRAMP, dbAMP);
- **Average identity**: $<35\%$ to training data AMPs;
- **Structural diversity**: Spanning $\alpha$-helix-rich, $\beta$-sheet-rich, and mixed architectures.

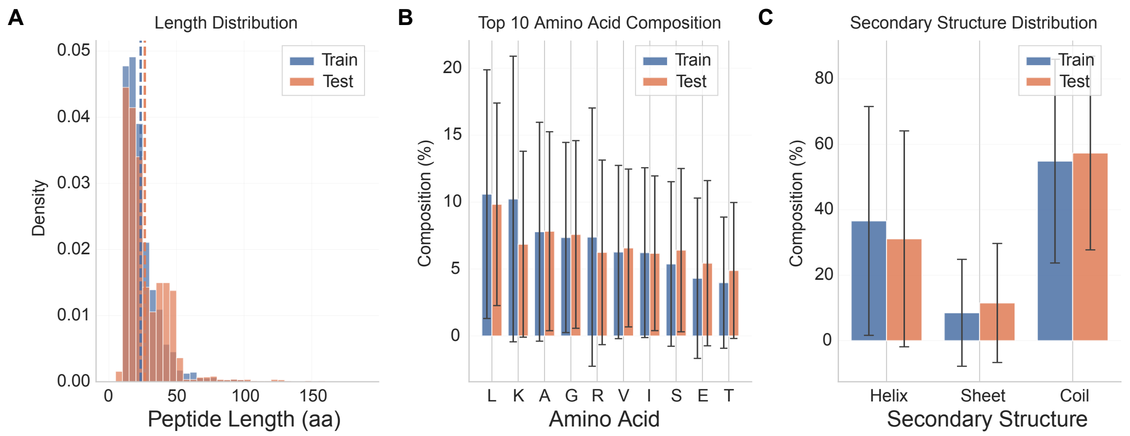

**Supplementary Figure 1: Comprehensive distribution comparison between training and test sets.** This figure provides a detailed comparison of peptide characteristics between the training and test datasets across three key dimensions: sequence length, amino acid composition, and secondary structure content. **(A)** shows the length distribution, with training set averaging 23.5 ± 13.6 amino acids (range: 11-190) and test set averaging 26.6 ± 17.2 amino acids (range: 5-183). **(B)** displays the amino acid composition comparison, revealing high compositional similarity between training and test sets across all 20 amino acids. **(C)** presents the secondary structure distribution comparison, showing similar structural profiles with test set exhibiting slightly higher helix content (37.9% vs 35.2%) and training set showing marginally more coil regions (47.8% vs 45.1%). The overall distributional consistency between training and test sets ensures fair evaluation while the low sequence similarity (Figure 2A in main text) guarantees robust generalization assessment.

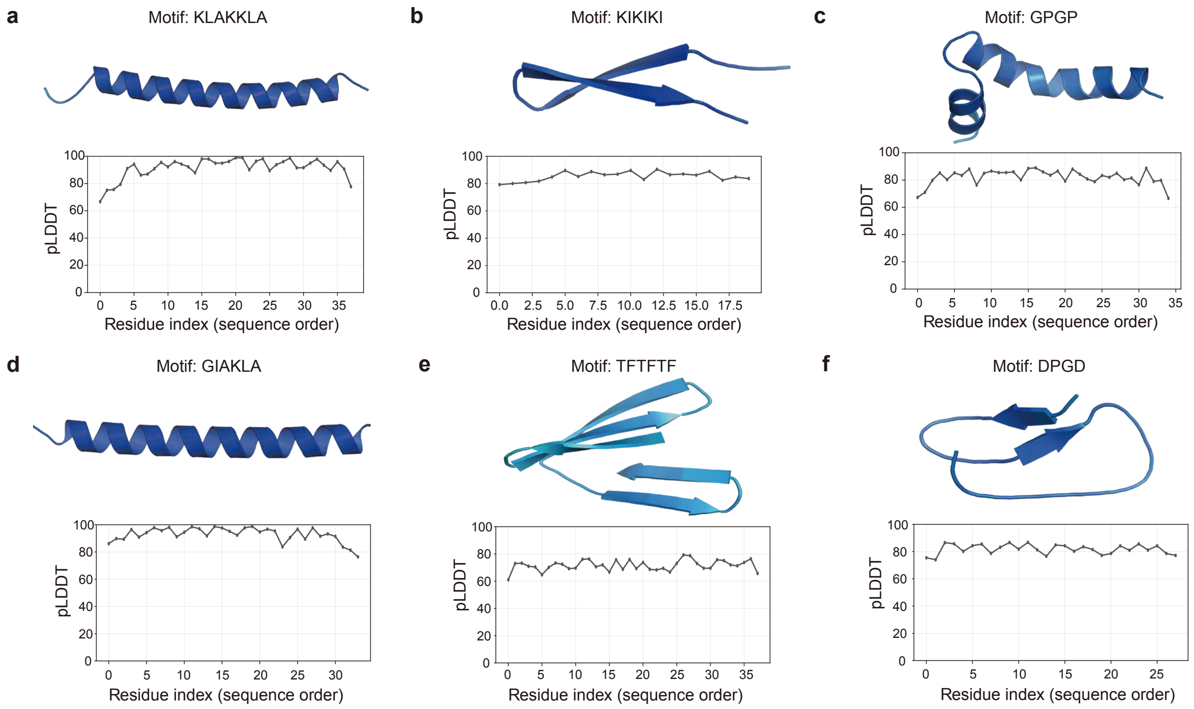

**Supplementary Figure 2: Predicted structures and confidence scores for representative protein motifs.** The figure displays the predicted three-dimensional structures and corresponding per-residue confidence scores (pLDDT) for six distinct protein motifs. For each panel, the top image shows the predicted structure as a ribbon diagram coloured by pLDDT value, while the bottom plot shows the pLDDT score for each residue along the sequence. The panels show: **a**, an amphipathic $\alpha$-helix motif (magainin-like), KLAKKLA; **b**, an alternating charged-hydrophobic $\beta$-strand, KIKIKI; **c**, a proline-glycine turn, GPGP; **d**, a short amphipathic helix, GIAKLA; **e**, a threonine-phenylalanine $\beta$-strand, TFTFTF; and **f**, a charged turn motif, DPGD. (All results here have an amp score greater than 0.95)
